# Breast invasive ductal carcinoma classification on whole slide images with weakly-supervised and transfer learning

**DOI:** 10.1101/2021.07.06.451320

**Authors:** Fahdi Kanavati, Masayuki Tsuneki

## Abstract

Invasive ductal carcinoma (IDC) is the most common form of breast cancer. For the non-operative diagnosis of breast carcinoma, core needle biopsy has been widely used in recent years which allows evaluation of both cytologic and tissue architectural features; so that it can provide a definitive diagnosis between IDC and benign lesion (e.g., fibroadenoma). Histopathological diagnosis based on core needle biopsy specimens is currently the cost effective method; therefore, it is an area that could benefit from AI-based tools to aid pathologists in their pathological diagnosis workflows. In this paper, we trained an Invasive Ductal Carcinoma (IDC) Whole Slide Image (WSI) classification model using transfer learning and weakly-supervised learning. We evaluated the model on a core needle biopsy (n=522) test set as well as three surgical test sets (n=1129) obtaining ROC AUCs in the range of 0.95-0.98.

## Introduction

Breast cancer is one of the leading causes of global cancer incidence [1]. In 2020, there were 2,261,419 new cases (11.7% of all cancer cases) and 684,996 deaths (6.9% of all cancer related deaths) due to breast cancer. Among women, breast cancer accounts for 1 in 4 cancer cases and for 1 in 6 cancer deaths in the vast majority of countries (159 of 185 countries) [1].

Invasive Ductal Carcinoma (IDC) (or invasive carcinoma of no special type: ductal NST) is a heterogenous group of tumors which fail to exhibit sufficient characteristics to achieve classification as a specific histopathological types. Microscopically, there are a wide variety of histopathological characteristics in IDCs. IDC grows in diffuse-sheets, well-defined nests, cords, or as individual (single) cells. Tubular differentiation tends to be well developed, barely detectable, or altogether absent.

Core needle biopsy is frequently used for the management of the non-palpable mammogram abnormalities, as it is cost effective and provides an alternative to short-interval follow-up mammography. It is also generally favored over fine-needle aspiration biopsy (FNAB) for the non-operative diagnosis of breast carcinoma, and it could replace open breast biopsy provided that the quality assurance is acceptable [2, 3]. Core needle biopsy allows the evaluation of both cytopathological and histopathological features, making it possible to provide a definitive diagnosis of IDC and benign lesions (e.g., fibroadenoma) in over 90% of cases [4]. All these factors highlight the benefit of establishing a histopathological screening system based on core needle biopsy specimens for breast IDC patients. Glass slides of biopsy specimens can be digitised as Whole Slide Images (WSIs) and could benefit from the application of computational histopathology algorithms to aid pathologists as part of a screening system.

Deep learning has found a wide array of applications in computational histopathology in the past few years. The applications from cancer cells classification and segmentation and patient outcome predictions for a variety of organs and diseases [5, 6, 7, 8, 9, 10, 11, 12, 13, 14, 15, 16, 17, 18]. Machine learning has been previously applied to various applications of breast histopathology classification [19, 20, 21, 22, 23, 24].

In this paper, we trained a WSI breast IDC classification model using transfer learning from ImageNet and weakly-supervised learning. We have also evaluated on the test sets, without fine-tuning, models that had been previously trained on other organs for the classification of carcinomas.

## Results

### A deep learning model for WSI breast IDC classification

The purpose of this study was to train a deep learning model to classify breast IDC in WSIs. We had a total of 1,154 biopsy WSIs of which we used 632 for training and 522 for testing. In addition, we used 1,129 surgical WSIs obtained from three sources as part of supplementary test sets. We used a transfer learning approach based on partial fine-tuning [25] to train the models. Figure 1 shows an overview of our training method. We then evaluated the trained models on four tests sets: one biopsy test set and three surgical test sets. As we had at our disposal six models [18, 26, 27, 28, 29, 30] that had been trained specifically on specimens from different organs (stomach, colon, lung, and pancreas), we evaluated those models without fine-tuning on the test sets to investigate whether morphological cancer similarities transfer across organs without additional training. Table 2 breaks down the distribution of the WSIs in each test set. For each test set, we computed the ROC AUC and log loss, and we have summarised the results in Tab. 1 and Fig. 2. Figures 4, 5, and 7, show representative heatmap prediction outputs for true positive, false positive, and false negative. Table 3 shows a confusion matrix breakdown by subtype for the false positives and true negatives using a probability threshold of 0.5. All 10 false positive WSIs were fibroadenomas. Figure 6 shows overview of representative fibroadenoma histopathology of 10 cases (WSIs) which were falsely predicted as IDC. There were representative histopathologic changes (e.g., proliferative epithelial changes, fibrocystic epithelial canges, and stromal changes) [31] in falsely predicted fibroadenomas (Figure 6); the proliferative findings could be the potential cause of the false positive.

**Figure 1:**
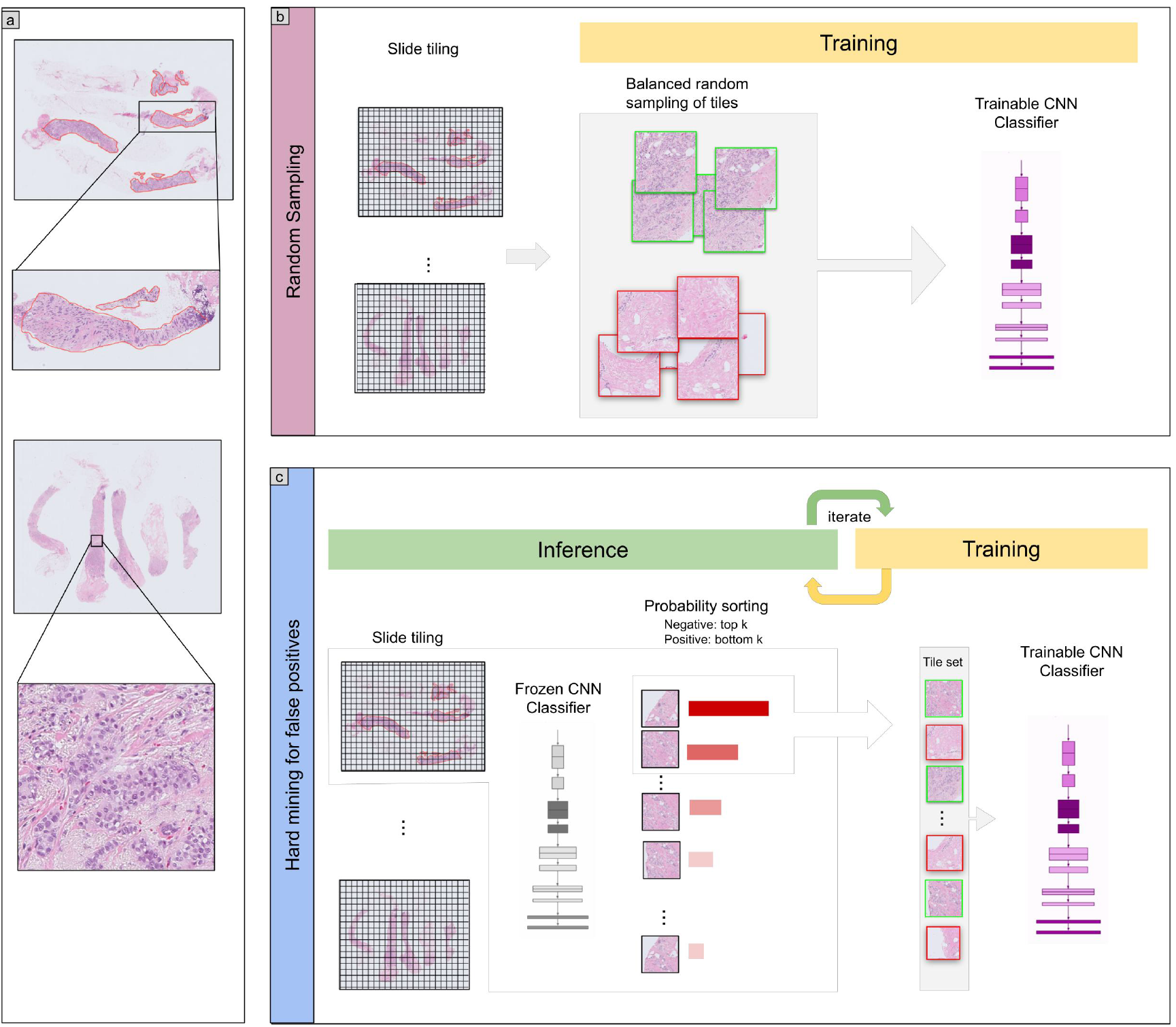
Overview. Training consisted of two stages. In the first stage (b) we randomly sampled tiles from the positive and negative WSIs, restricting the sampling from WSIs that had annotations if they were positive. In the second stage (c) We iteratively alternated between inference and training, relying only on the WSI label. During inference, the model weights were frozen, and it was applied in a sliding window fashion on each WSI. The top k tiles with the highest probabilities were then selected from each WSI. During training the selected tiles were then used to train the model.

**Table 1:**
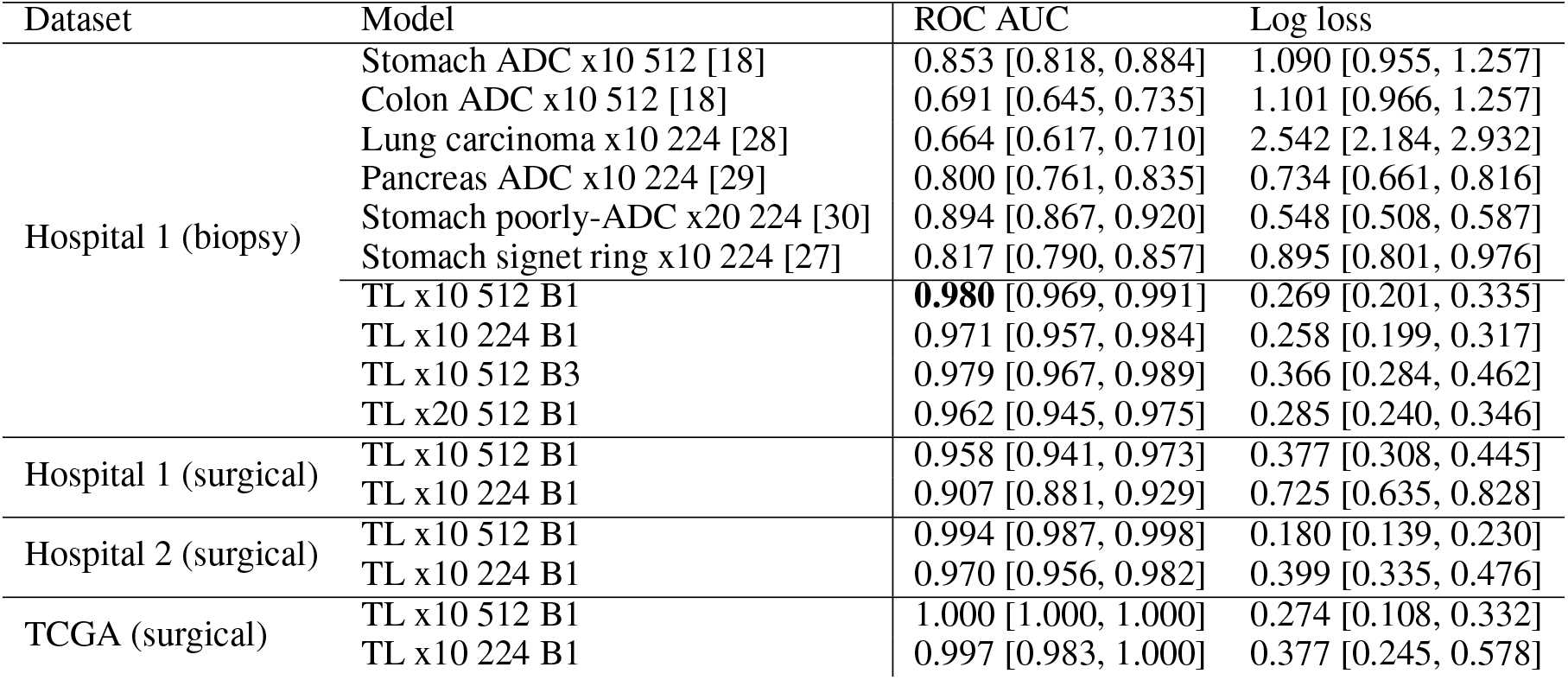
ROC and log loss results of the models on the biopsy and surgical test sets

**Table 2:**
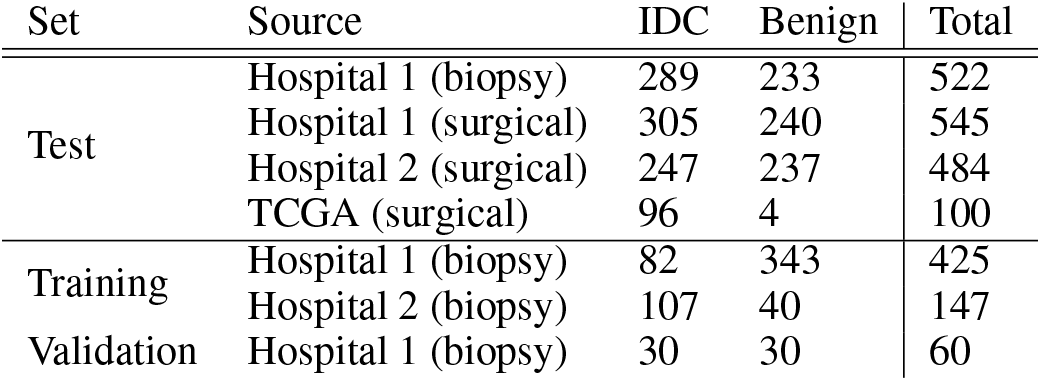
Distribution of WSIs in the different sets.

| Set | Source | IDC | Benign | Total |
| --- | --- | --- | --- | --- |
| Test | Hospital 1 (biopsy) | 289 | 233 | 522 |
|  | Hospital 1 (surgical) | 305 | 240 | 545 |
|  | Hospital 2 (surgical) | 247 | 237 | 484 |
|  | TCGA (surgical) | 96 | 4 | 100 |
| Training | Hospital 1 (biopsy) | 82 | 343 | 425 |
|  | Hospital 2 (biopsy) | 107 | 40 | 147 |
| Validation | Hospital 1 (biopsy) | 30 | 30 | 60 |

**Figure 2:**
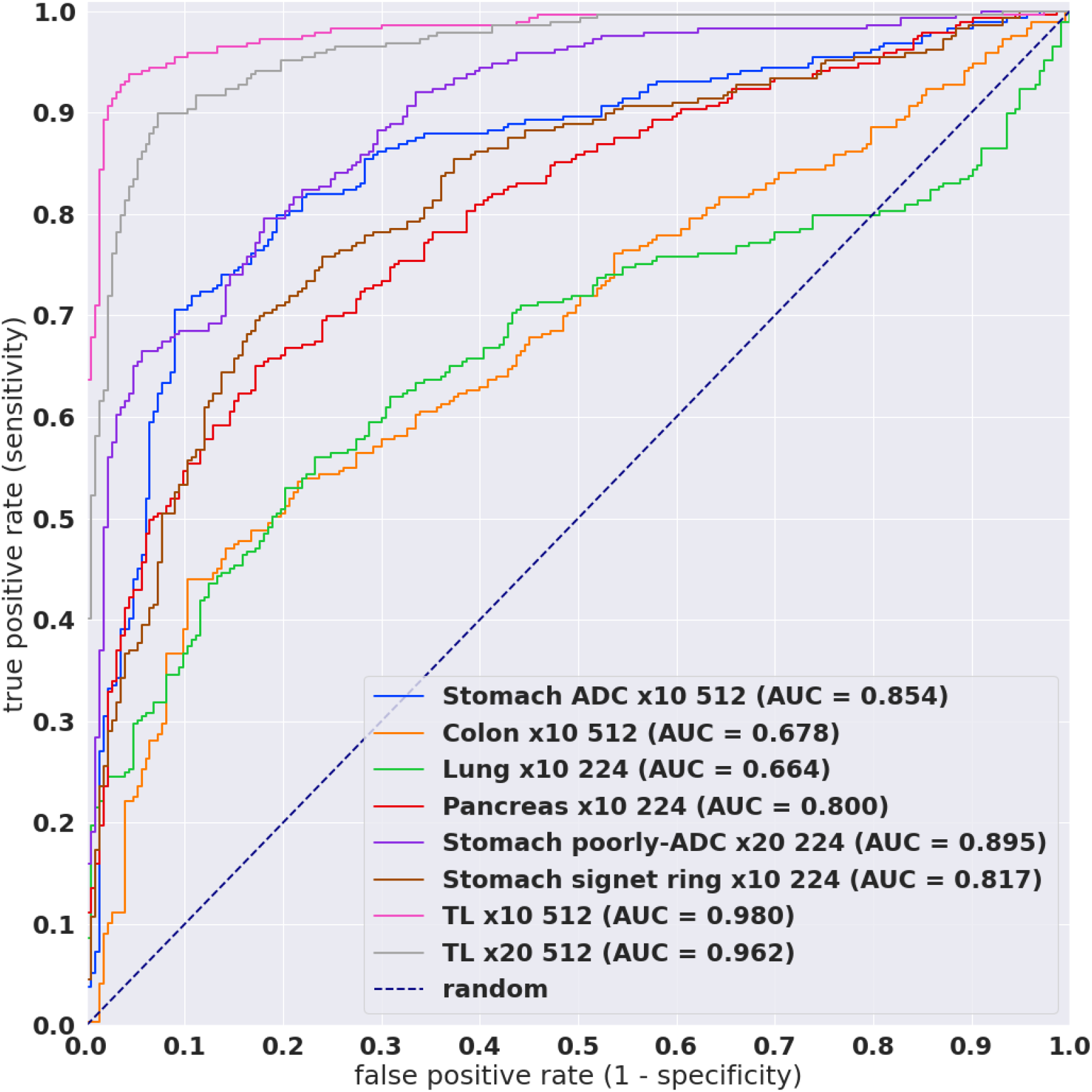
ROC curves for the various existing models as well as models trained via transfer learning on core needle biopsy test set.

**Figure 3:**
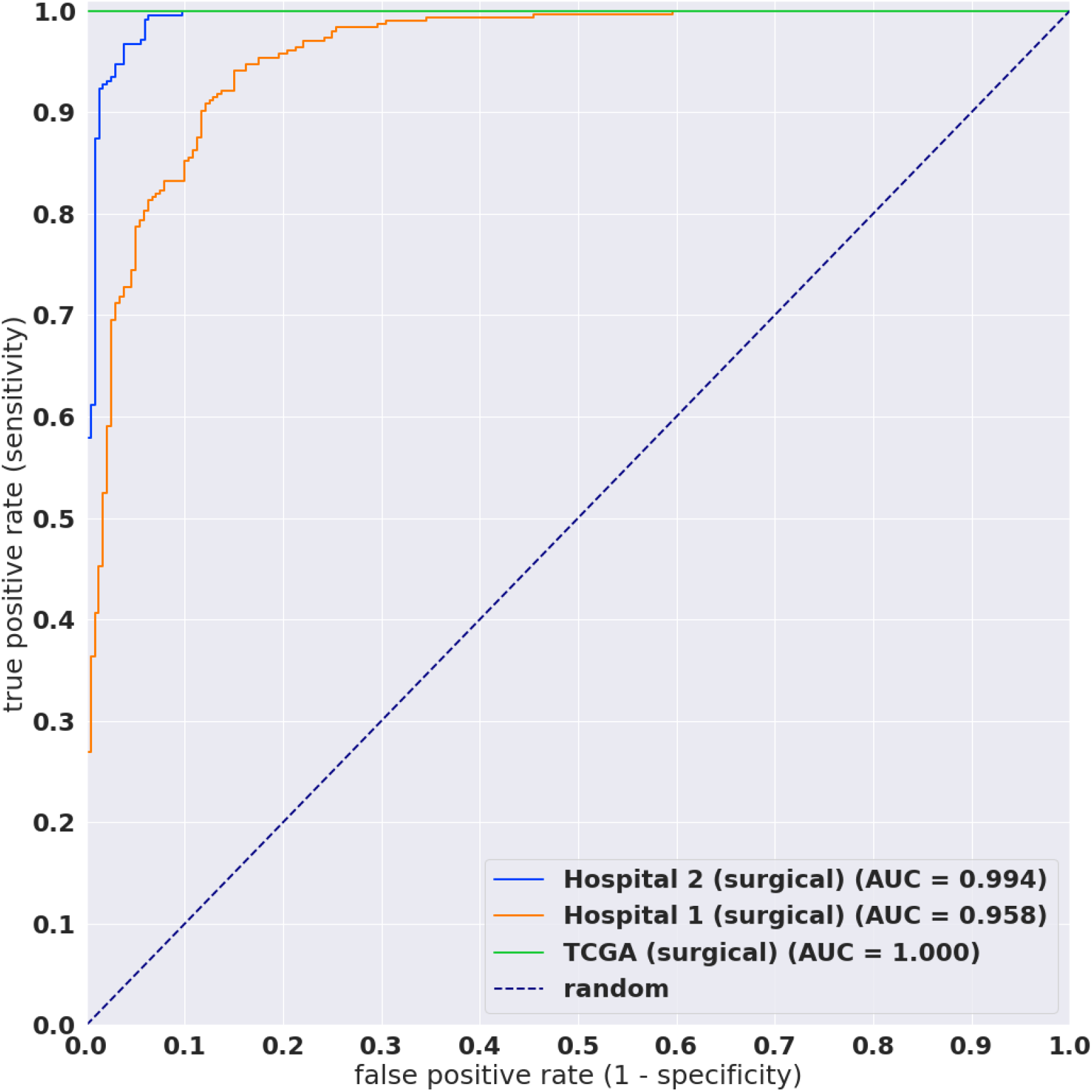
ROC curves on surgical test sets.

**Figure 4:**
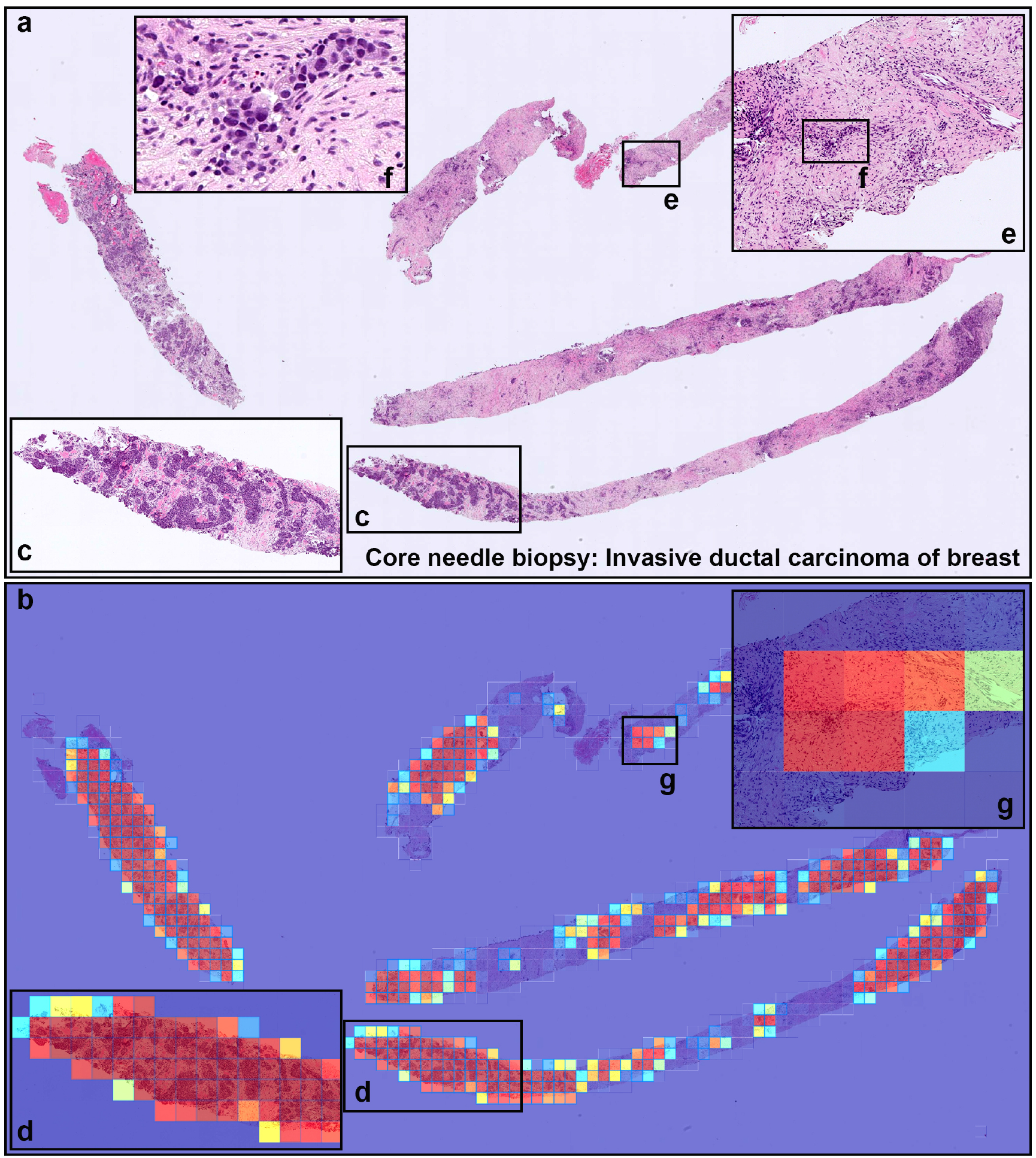
A representative true positive invasive ductal carcinoma (IDC) of breast from core needle biopsy test set. Heatmap images show true positive predictions of IDC cells (b) and they correspond respectively to H&E histopathology (a) using transfer learning from ImageNet model (magnification x10). Not only abundant IDC cells invading areas (c) but also a few IDC cells (e, f), heatmap images show appropriately true positive predictions (d, g).

**Figure 5:**
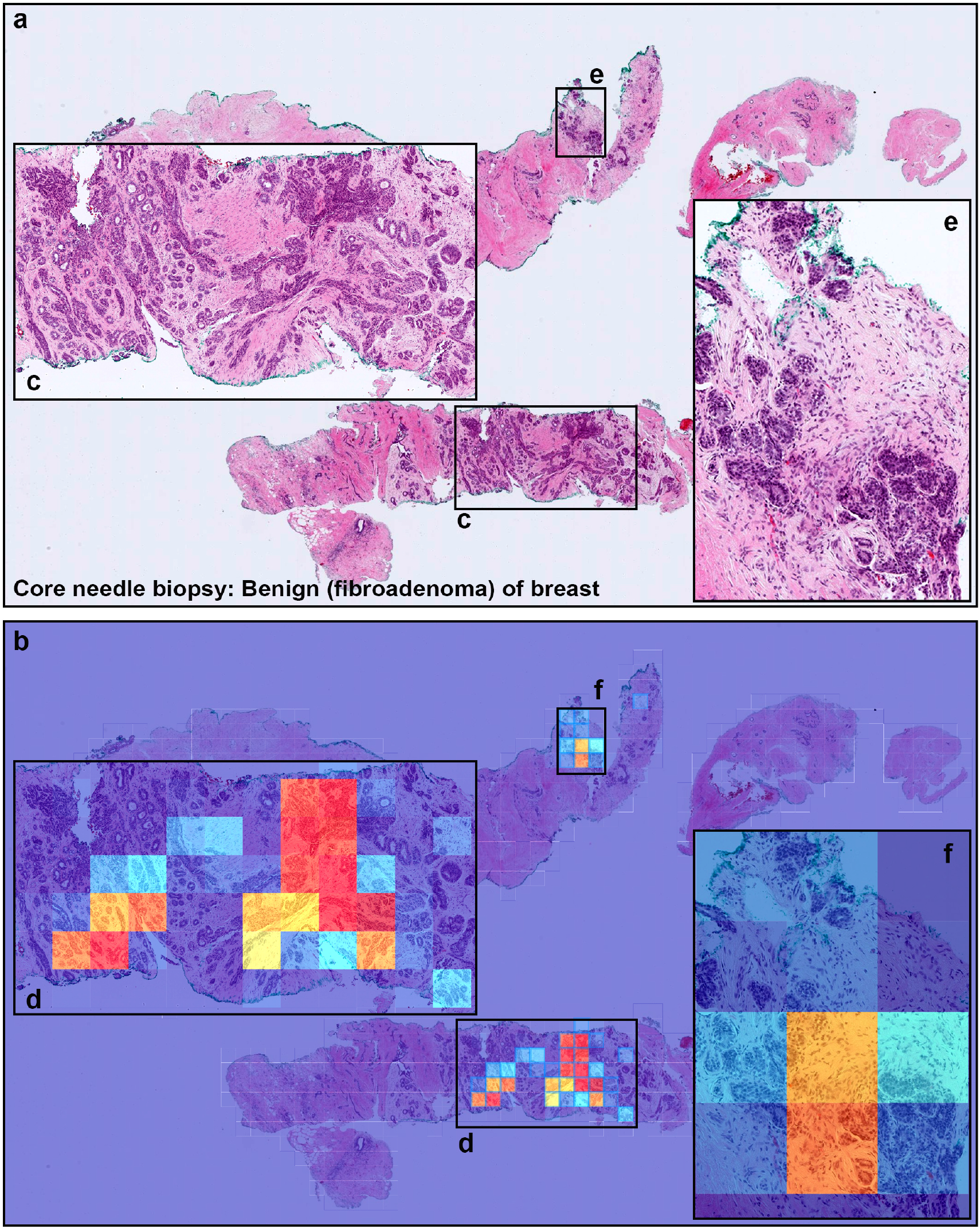
A representative example of invasive ductal carcinoma (IDC) false positive prediction output on a case from core needle biopsy test set. Histopathologically (a), this case is a benign lesion (fibroadenoma). Heatmap images (b, d, exhibited false positive prediction of IDC using transfer learning from ImageNet model (magnification x10). The ductular structures in fibroadenoma with a pericanalicular pattern (c, d, e, f) would be the primary cause of false positive due to its morphological analogous to ductular structures in IDC.

**Figure 6:**
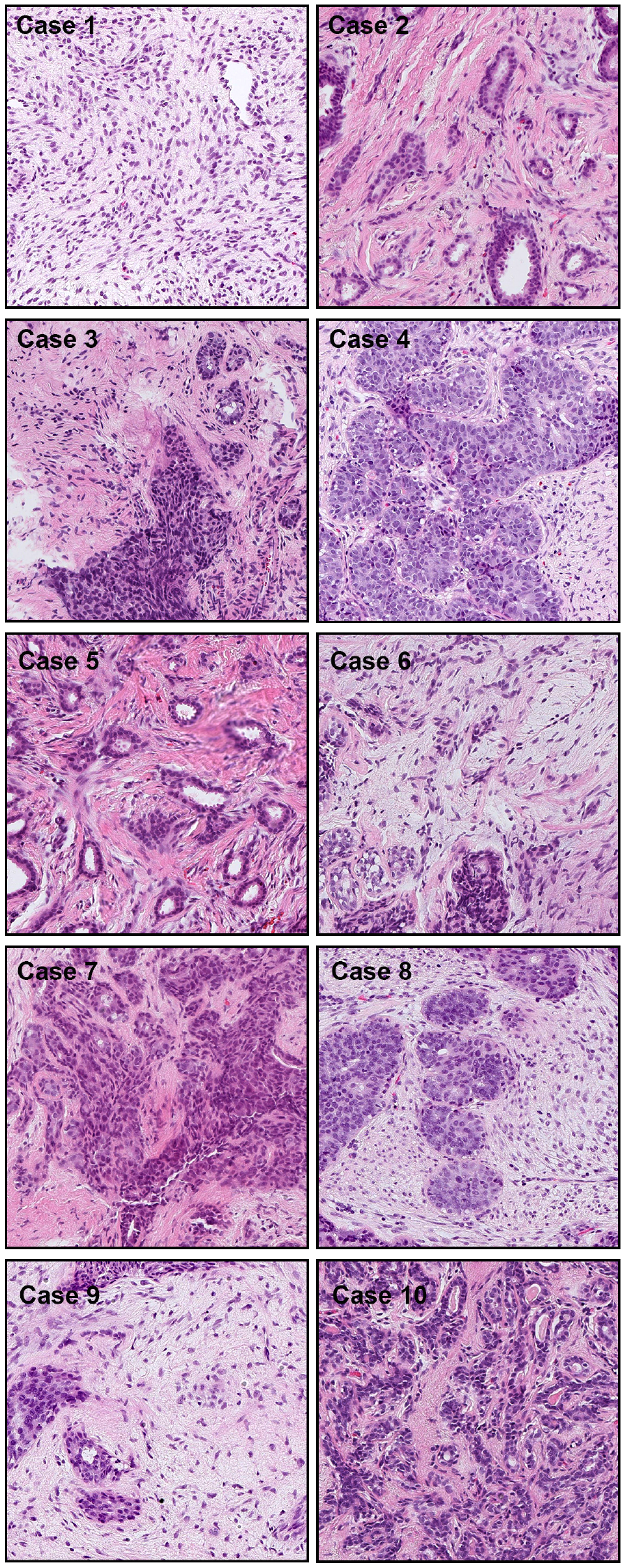
Representative tissue areas (Cases 1-10), without heatmap overlay, that were falsely predicted as IDC. There were 10 cases of false positive prediction outputs from the core needle biopsy test set. The false positive predictions are most likely due to the enlarged spindle shaped stromal cell nuclei with pleomorphism and tubules composed of cuboidal or low columnar cells with round uniform nuclei resting on a myoepithelial cell layer. This is morphologically analogous to invading single cells, ductular structures, and cancer stroma in IDC.

**Figure 7:**
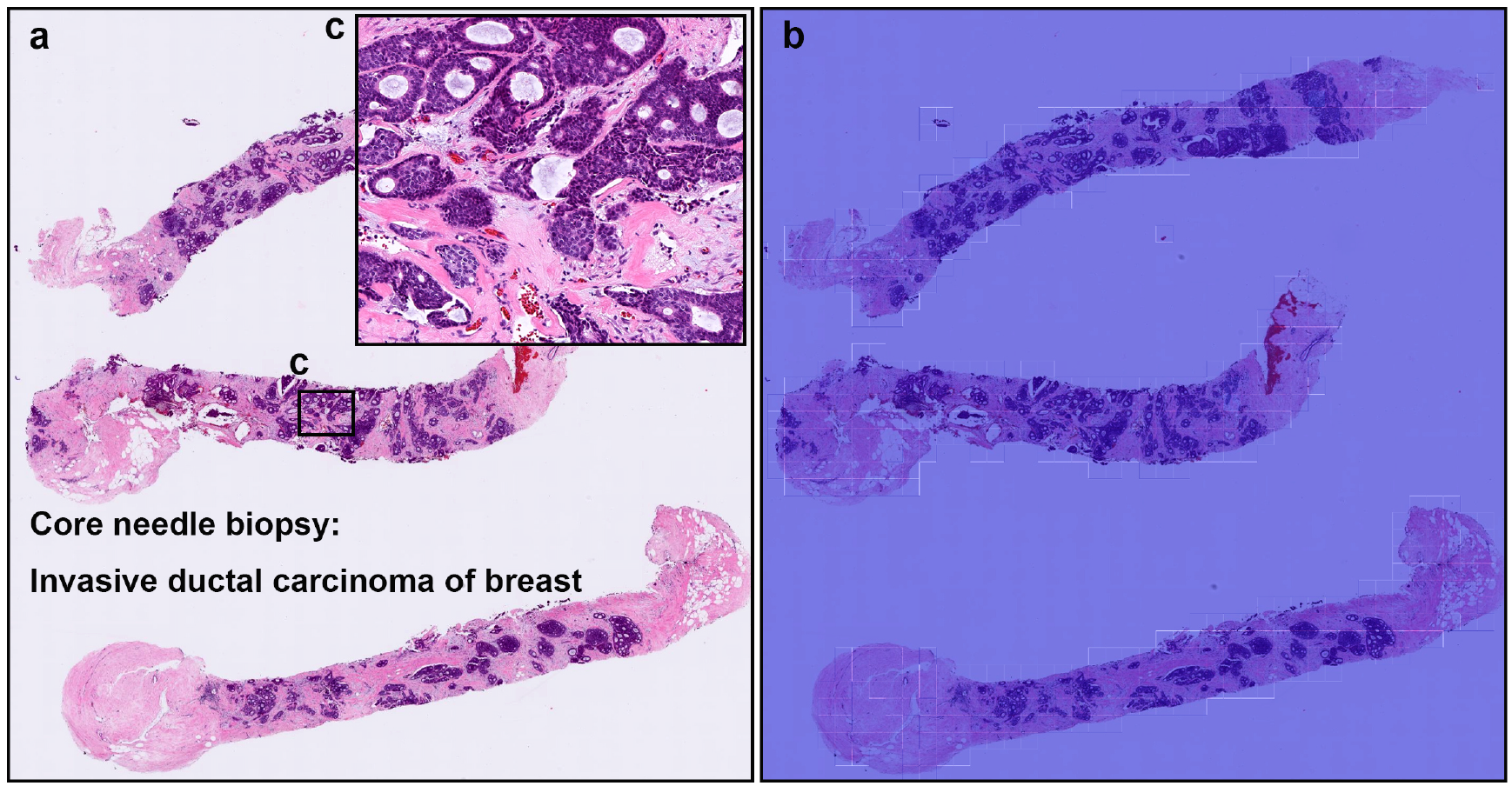
A representative false negative prediction output on a case from core needle biopsy test set. According to the histopathological report, this case (a, c) is an invasive ductal carcinoma (IDC). However, there is no true positive predictions of IDC cells on heatmap image (b).

**Figure 8:**
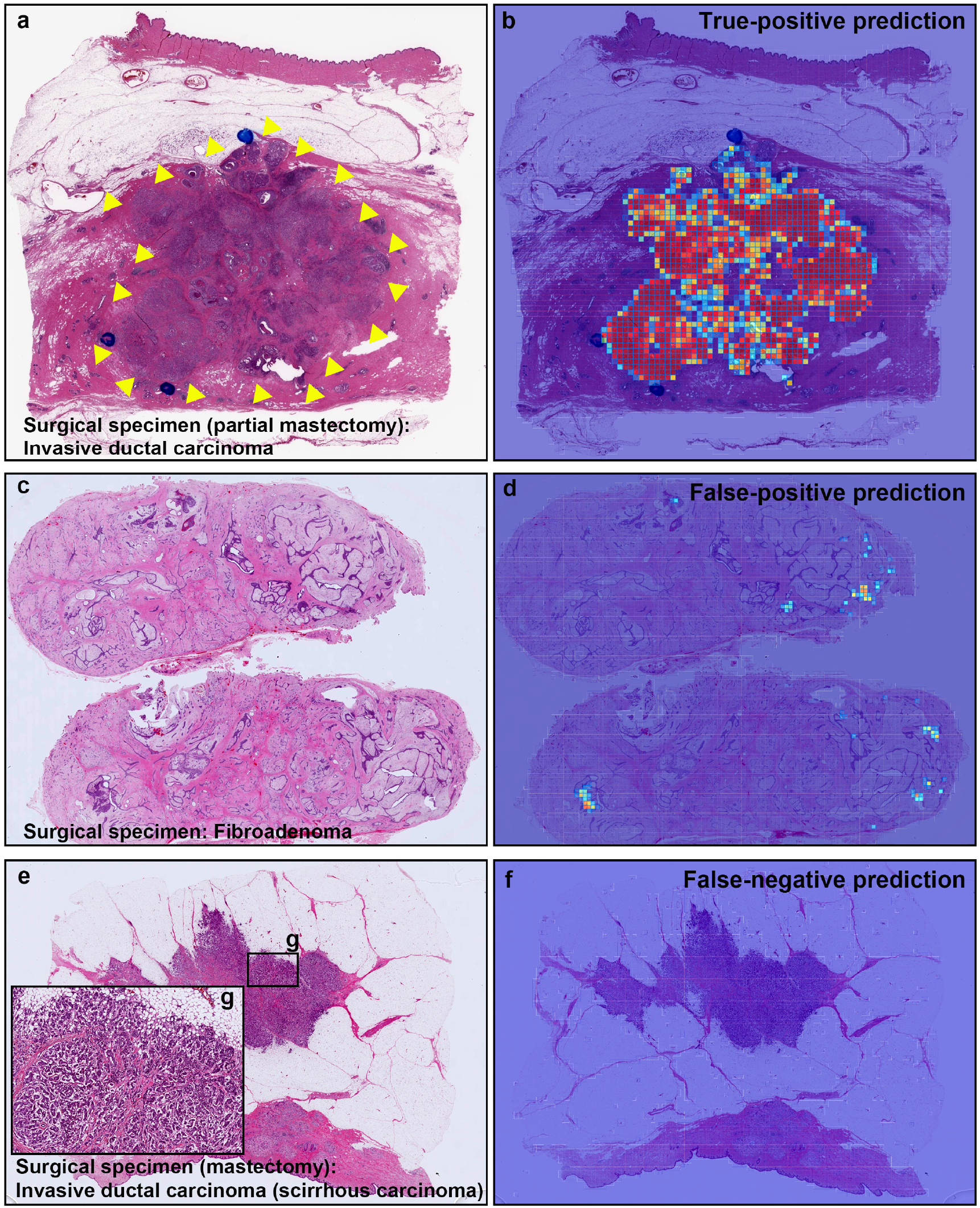
Representative true positive, false positive, and false negative prediction outputs on surgically resected specimens for invasive ductal carcinomas (IDCs) and fibroadenoma. Histopathologically, (a) has IDC; (c) is fibroadenoma; and (e) has IDC (scirrhous type). (b) shows true positive probability heatmap using transfer learning from ImageNet model (magnification x10) for IDC invading area which was corresponded to surgical pathologists marked area with blue-ink-dots (and yellow-triangles) (a). (d) exhibited false positive prediction of IDC in fibroadenoma. There is no true positive predictions of IDC cells on heatmap image (f) in scirrhous carcinoma of IDC (e).

**Table 3:** A breakdown of the subtypes of the false positives and true negatives in the biopsy test set using the TL model x10 using a classification threshold of 0.5.

|  | Subtype of benign | number of WSIs | % |
| --- | --- | --- | --- |
| False-positives (10 WSIs) | Fibroadenoma | 10 | 100.0 |
| True-negatives (223 WSIs) | Fibroadenoma | 121 | 54.3 |
|  | Mastopathy | 67 | 30.0 |
|  | Normal | 24 | 10.8 |
|  | Fibrosis | 6 | 2.7 |
|  | Ductal hyperplasia | 2 | 0.9 |
|  | Granulation tissue | 2 | 0.9 |
|  | Fat necrosis | 1 | 0.4 |

## Discussion

In this study, we trained deep learning models for the classification of breast IDC in surgical and biopsy WSIs. We used weakly-supervised and transfer learning. We used the partial fine-tuning approach, which is fast to train. The best model achieved AUCs in the range of 0.96-0.98.

Overall, the EfficientNetB1 model trained at magnification x10 achieved slightly better results than x20 on the biopsy test set. In addition, using a larger tile size of 512×512px achieved slightly better results than 224×244px. Despite IDC morphology having some similarity with adenocarcinoma, the application of models that classify ADC on other organs did not fully generalise to IDC. The stomach ADC had the highest AUC 0.85-0.89 when applied to breast IDC WSIs. While the results on the TCGA test set are high, it does not provide a proper evaluation in terms of potential false positives as there were only four benign cases.

All of the false positive cases in the biopsy test set were fibroadenomas (see Tab. 3). Fibroadenomas exhibit a wide range of morphology, and it could be that the variety was not fully represented in the training set, which only had 91 cases of fibroadenomas compared to the 131 in the test set. One source of difficulty in creating a balanced set of the fibroadenomas varieties is that the diagnostic reports did not include detailed description of fibroadenoma histology, making a simple random partition the only option. In addition, the test set had a larger proportion of fibroadenomas compared to other benign subtypes. Therefore, as future work, it would be important to investigate the histopathological typing of fibroadenomas in order to develop better deep learning models.

According to the guideline by General Rule Committee of the Japanese Breast Cancer Society [32], the pathological diagnosis of IDC is sufficient for core needle biopsy. Therefore, the application of a deep learning model, once properly validated, in a clinical setting would help pathologists in their diagnostic workflows. On the other hand, surgical specimens tend to require further subtyping of IDC, so future work could look into developing models specifically for IDC subtype classification for surgical specimens.

## Methods

### Clinical cases and pathological records

This is a retrospective study. A total of 2,183 H&E (hematoxylin & eosin) stained histopathological specimens of human breast IDC and benign lesions – 1,154 core needle biopsy and 1,028 surgical – were collected from the surgical pathology files of three hospitals: International University of Health and Welfare, Mita Hospital (Tokyo) and Kamachi Group Hospitals (Fukuoka) after histopathological review of those specimens by surgical pathologists. The test cases were selected randomly, so the obtained ratios reflected a real clinical scenario as much as possible. All WSIs were scanned at a magnification of x20. In addition, we collected 100 WSIs from TCGA; however, only four bengin cases were available.

### Dataset

The pathologists excluded cases that were inappropriate or of poor scanned quality prior to this study. The diagnosis of each WSI was verified by at least two pathologists. Table 2 breaks down the distribution of dataset into training, validation, and test sets. Hospitals which provided histopathological cases were anonymised (e.g., Hospital 1-2). The training set was solely composed of WSIs of core needle biopsy specimens. The test sets were composed of WSIs of core needle biopsy or surgical specimens. The patients’ pathological records were used to extract the WSIs’ pathological diagnoses and to assign WSI labels. 96 WSIs out of the 191 WSIs with IDC were loosely annotated by pathologists. The rest of IDC and benign WSIs were not annotated and the training algorithm only used the WSI labels. Each WSI diagnosis was observed by at least two pathologists, with the final checking and verification performed by a senior pathologist.

### Deep learning models

We trained all the models using the partial fine-tuning approach [25]. This method consists in using the weights of an existing pre-trained model and only fine-tuning the affine parameters of the batch normalisation layers and the final classification layer. We have used the EfficientNetB1 architecture [33], as well as B3, with a modified input sizes of 224×224px and 512×512px, starting with pre-trained weights from ImageNet. The total number of trainable parameters for EfficientNetB1 was only 63,329.

The training method that we have used in this study is exactly the same as reported in a previous study [26] with the main difference being the use of partial fine-tuning method. For completeness, we repeat the method here.

To apply the model on the WSIs for training and inference, we performed slide tiling by extracting fixed-sized tiles from tissue regions. We detected the tissue regions by performing a thresholding on a grayscale version of the WSI using Otsu’s method [34], which allows the elimination of most of the white background. During inference, we performed the slide tiling in a sliding window fashion on the tissue regions, using a fixed-size stride that was half the size of the tile. During training, we initially performed random balanced sampling of tiles from the tissue regions, where we maintained an equal balance of positive and negative labelled tiles in the training batch. To do so, we placed the WSIs in a shuffled queue with oversampling of the positive labels to ensure that all the WSIs were seen at least once during each epoch, and we looped over the labels in succession (i.e. we alternated between picking a WSI with a positive label and a negative label). Once a WSI was selected, we randomly sampled 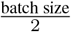 tiles from each WSI to form a balanced batch. We then switched into hard mining of tiles. To perform the hard mining, we alternated between training and inference. During inference, the CNN was applied in a sliding window fashion on all of the tissue regions in the WSI, and we then selected the *k* tiles with the highest probability for being positive. If the tile is from a negative WSI, this step effectively selects the false positives. The selected tiles were placed in a training subset, and once the subset size reached *N* tiles, a training pass was triggered. We used *k* = 4, *N* = 256, and a batch size of 32.

A subset of WSIs with IDC were loosely annotated (n=96) while the rest had WSI-level labels only (n=95). From the loosely annotated WSIs, we only sampled tiles from the annotated tissue regions. Otherwise, we freely sampled tiles from the entire tissue regions.

The models were trained on WSIs at x10 and x20 magnifications. We used two input tile sizes: 512×512px and 224×224px. The strides were half the tile sizes. The WSI prediction was obtained by taking the maximum probability from all of the tiles.

We trained the models with the Adam optimisation algorithm [35] with the following parameters: *beta*_1_ = 0.9, *beta*_2_ = 0.999. We used a learning rate of 0.001. We applied a learning rate decay of 0.95 every 2 epochs. We used the binary cross entropy loss. We used early stopping by tracking the performance of the model on a validation set, and training was stopped automatically when there was no further improvement on the validation loss for 10 epochs. The model with the lowest validation loss was chosen as the final model.

### Software and statistical analysis

The deep learning models were implemented and trained using TensorFlow[36]. AUCs were calculated in python using the scikit-learn package[37] and plotted using matplotlib [38]. The 95% CIs of the AUCs were estimated using the bootstrap method[39] with 1000 iterations.

## Data availability

Due to specific institutional requirements governing privacy protection, datasets used in this study are not publicly available.

